# An almost nontoxic tetrodotoxin analog, 5,6,11-trideoxytetrodotoxin, as an odorant for the grass puffer

**DOI:** 10.1101/2021.09.12.459881

**Authors:** Yoshihisa Noguchi, Takehisa Suzuki, Keigo Matsutani, Ryo Sakakibara, Ryota Nakahigashi, Masaatsu Adachi, Toshio Nishikawa, Hideki Abe

**Affiliations:** Laboratory of Fish Biology, Graduate School of Bioagricultural Sciences, Nagoya University, Furo-Cho, Chikusa-Ku, Nagoya, Aichi 464-8601, Japan; Laboratory of Organic Chemistry, Graduate School of Bioagricultural Sciences, Nagoya University, Furo-Cho, Chikusa-Ku, Nagoya, Aichi 464-8601, Japan

**Keywords:** Tetrodotoxin (TTX), 5,6,11-trideoxyTTX, toxification, odorant, pheromone

## Abstract

Toxic puffers contain the potent neurotoxin, tetrodotoxin (TTX). Although TTX is considered to serve as a defense substance, previous behavioral studies have demonstrated that TTX acts as an attractive pheromone for some toxic puffers. To elucidate the physiological mechanism of putative pheromonal action of TTX, we examined whether grass puffers *Takifugu alboplumbeus* can detect TTX. Electroolfactogram (EOG) results suggest that the olfactory epithelium (OE) of grass puffers responded to a type of TTX analog (5,6,11-trideoxyTTX), although it did not respond to TTX. We also examined the attractive action of 5,6,11-trideoxyTTX on grass puffers by recording their swimming behavior under dark conditions. Grass puffers preferred to stay on the side of the aquarium where 5,6,11-trideoxyTTX was administered, and their swimming speed decreased. Additionally, odorant-induced labeling of olfactory sensory neurons by immunohistochemistry against neural activity marker (phosphorylated extracellular signal regulated kinase; pERK) revealed that labeled olfactory sensory neurons were localized in the region surrounding “islets” where there was considered as nonsensory epithelium. 5,6,11-trideoxyTTX has been known to accumulate in grass puffers, but its toxicity is much lower (almost nontoxic) than TTX. Our results suggest that toxic puffers may positively use this TTX analog, which has been present in their body with TTX but whose function was unknown, as an odorant for chemical communication or effective TTX accumulation.

## Introduction

Toxic puffers accumulate the potent neurotoxin, tetrodotoxin (TTX), which is a well-known voltage-gated sodium channel blocker ^1^. TTX was first believed to be present only in pufferfish from the family Tetraodontidae. However, it was later detected in taxonomically diverse animals (e.g., teleosts, amphibians, echinoderms, crustaceans, mollusks, and worms) inhabiting terrestrial, marine, freshwater, and brackish water environments ^2^.

Regardless of the abundant studies, the biological origin of TTX and its ecological functions remain unclear. However, the presence of TTX in such a variety of organisms suggests that the ultimate origin of TTX is exogenous, produced by certain aquatic bacteria ^1,3^. TTX and its analogs were detected in pufferfish prey, including starfish, gastropods, crustaceans, flatworms, and ribbonworms, in addition to several species of bacteria that are free-living or symbiotic with pufferfish or their food organisms ^2^. Additionally, juveniles of the cultivated tiger puffer *Takifugu rubripes* became nontoxic in aquaria or cages that were suspended above the seafloor with a TTX-free diet ^3,4^. These results suggest that TTX in pufferfish is a result of its accumulation through the food chain. Such a phenomenon of a nontoxic organism becoming toxic by taking toxins into its body is called toxification.

Why do toxic puffers accumulate TTX in their bodies? Some toxic puffers have TTX-secreting glands on the skin ^5^, and electrical stimulation induced TTX secretion from the body surface of these puffers ^6^. In addition, TTX induces repulsive or avoidance reactions in predatory fish ^7^; predators that ingested *Takifugu* larvae spat them out promptly ^8,9^. Thus, in toxic puffers, TTX functions as a biological defense substance against predators.

Although the main function of TTX in pufferfish appears to be defensive, several studies have reported that pufferfish can exploit TTX as a source of communication. Sexually active male grass puffers (*T. alboplumbeus*) were attracted to crude TTX that was extracted from other pufferfish ^10^. Artificially raised nontoxic tiger puffer juveniles were also attracted to crude extract of TTX ^11–13^. These studies suggest that pufferfish may sense TTX. However, the TTX-sensing mechanism in these puffers remains unclear, although the attractive effect of TTX on tiger puffer juveniles disappeared after nose ablation ^11^.

In this study, we found that an almost nontoxic TTX analog, 5,6,11-trideoxyTTX (**Fig. 1**), that has also been known to accumulate in toxic puffers, is an odorant for the grass puffer. We demonstrated the presence of the olfactory sensing ability, specific olfactory sensory neurons (OSNs), and the chemotactic behavior of grass puffers in response to 5,6,11-trideoxyTTX. To the best of our knowledge, this is the first report that pufferfish positively use TTX analog, which has been present in their body with TTX but whose function was unknown, as an odorant.

**Figure 1:**
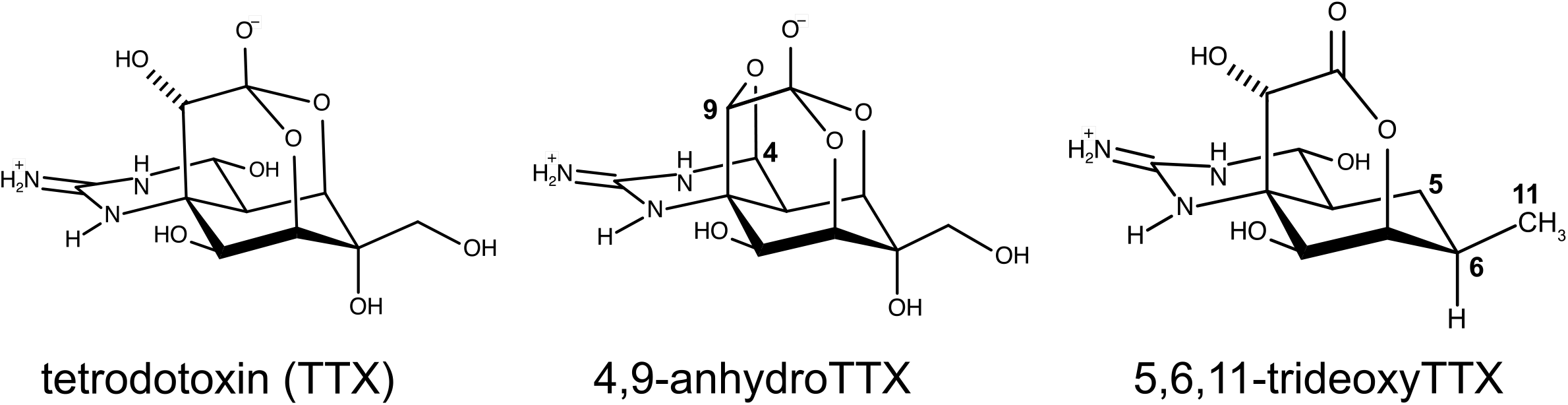
Molecular structures of tetrodotoxin and its analogs used in this study.

## Results

### EOG response of grass puffers to TTX

We first performed EOG recordings of adult grass puffers in response to commercially available TTX. The EOG response to a 10-s stimulus with an amino acid solution (10^−5^ M L-Arginine (L-Arg)) was a slow monophasic potential with a steep rising phase followed by an exponential decay toward the baseline (**Fig.2a**, bottom trace). However, TTX (10^−7^ and 10^−5^ M) did not induce significant changes in the EOG (**Fig. 2a** upper and middle traces, respectively). The EOG amplitude of 10^−7^ M TTX (0.027 ± 0.007 mV, n = 4; **Fig. 2b○**) and 10^−5^ M TTX (0.026 ± 0.007 mV, n = 7; **Fig. 2b●**) were significantly lower than that of 10^−5^ M L-Arg (0.074 ± 0.008 mV, n = 11; **Fig. 2b■**).

**Figure 2:**
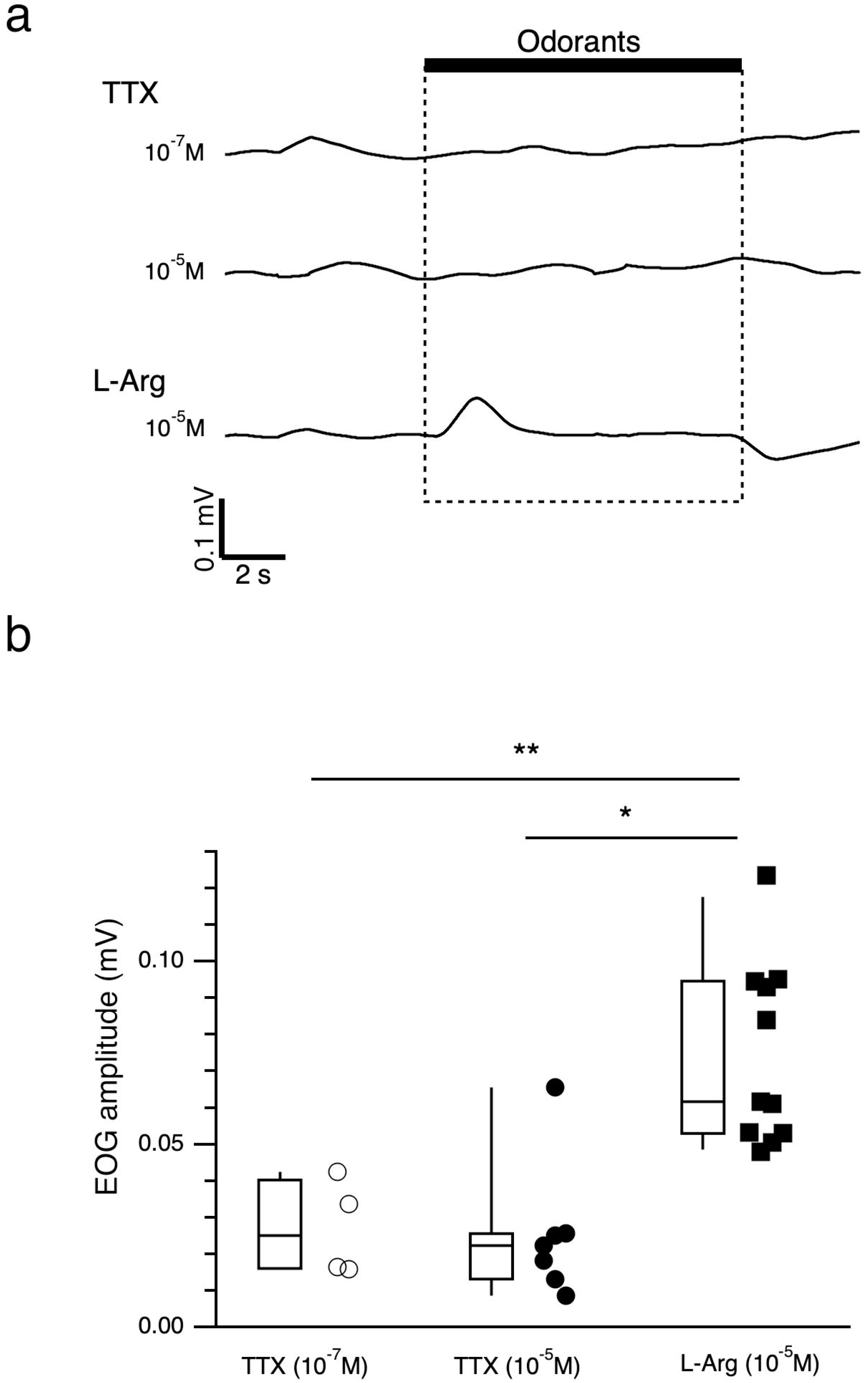
Grass puffer did not respond to TTX. **a**: EOG responses of the olfactory epithelium of *T. alboplumbeus* to TTX (10^−7^ M, upper trace; 10^−5^ M, middle trace) and L-Arg (10^−5^ M, under trace). **b**: EOG amplitudes of TTX (10^−7^ M, n = 6, open circle; 10^−5^ M, n = 7, filled circle) and L-Arg (10^−5^ M, n = 11, filled square). Box plots show the median, quartiles (boxes), and 10%–90% range (whiskers), and dot plots show raw EOG amplitudes. Steel’s multiple comparison test compared with L-Arg (TTX 10^−7^M, P = 0.009; TTX 10^−5^M, P = 0.011: * p<0.05, ** p<0.01.

### EOG response to TTX analogs

We then examined the EOG responses of the grass puffer to the two synthesized TTX analogs that have been reported to be simultaneously detected with TTX in toxic puffers or their prey’s bodies. A TTX analog, 4,9-anhydroTTX, did not induce significant changes in EOG traces (10^−5^ M; **Fig. 3a**, second trace) when the same concentration of L-Arg showed clear EOG responses (**Fig. 3a**, bottom trace). However, the same concentration of another TTX analog, 5,6,11-trideoxyTTX, induced a clear increase in EOG response (**Fig. 3a**, third trace). Vehicle control (2.5 × 10^−5^ N acetic acid solution in artificial seawater (ASW)) did not induce any changes in EOG traces (**Fig. 3a**, top trace). EOG amplitudes of 4,9-anhydroTTX, 5,6,11-trideoxyTTX, and L-Arg (all 10^−5^ M) were 0.027 ± 0.004 mV [n = 4], 0.197 ± 0.029 mV [n = 7], and 0.128 ± 0.017 mV [n = 7], respectively. EOG amplitudes of 5,6,11-trideoxyTTX and L-Arg were significantly larger than those of the vehicle (0.032 ± 0.007 mV [n = 5]; **Fig.3b**), and EOG amplitude of 5,6,11-trideoxyTTX showed a dose-dependent relationship (**Fig. 3c and d**). These results suggest that the OE of the grass puffers can detect 5,6,11-trideoxyTTX.

**Figure 3:**
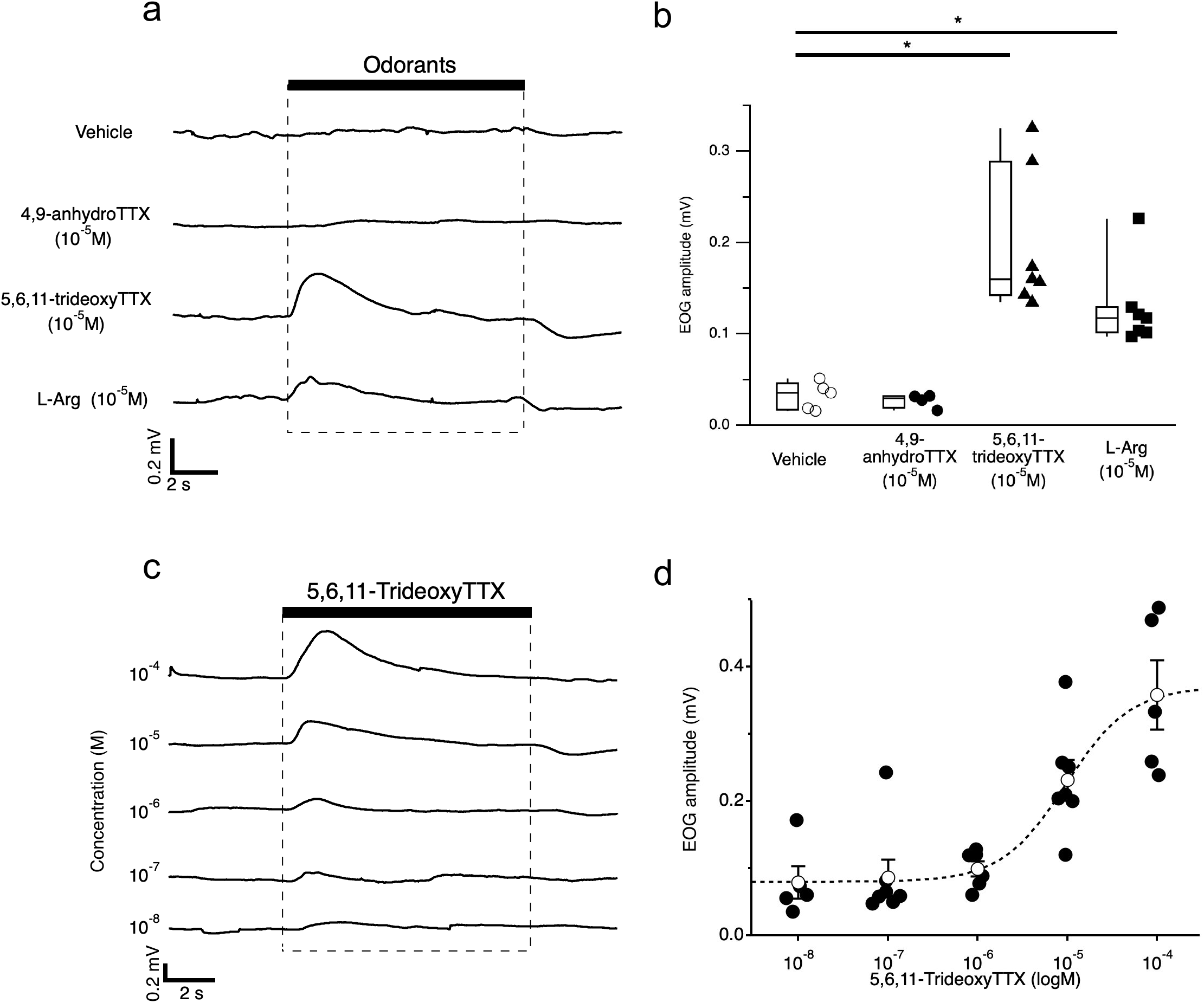
5,6,11-trideoxyTTX induced EOG response of the grass puffer. **a**: EOG responses to Vehicle (top trace), 4,9-anhydroTTX (10^−5^ M, second trace), 5,6,11-trideoxyTTX (10^−5^ M, third trace), and L-Arg (10^−5^ M, bottom trace) from same grass puffer. **b**: EOG amplitudes of Vehicle (n = 5, open circles), 4,9-anhydroTTX (n = 4, filled circles), 5,6,11-trideoxyTTX (n = 7, filled triangles), and L-Arg (n = 7, filled square). Box plots show the median, quartiles (boxes), and 10%–90% range (whiskers) of EOG amplitudes, and dot plots show the raw value of EOG amplitudes. Steel’s multiple comparison test compared with Vehicle (5,6,11-trideoxyTTX, P = 0.012; L-Arg, P = 0.012): * p < 0.05. **c:** Representative EOG responses by a grass puffer were evoked by increasing the concentration of 5,6,11-trideoxyTTX. The period of 5,6,11-trideoxyTTX administration is shown by the black bar. **d:** Dose-response relationship of 5,6,11-trideoxyTTX obtained from n = 5–7 fish. Peak EOG responses are plotted as a function of the concentrations delivered. Black circles show the individual EOG amplitudes, and open circles show the mean EOG amplitude at given concentrations. Error bars represent the standard error. The dotted line represents a Hill equation fitted to the data (EC_50_: 9.62 × 10^−6^ M, Hill coefficient: 1.21).

### Behavioral experiments

We next examined the behavioral response of adult grass puffers to 5,6,11-trideoxyTTX. One mature grass puffer was put in a test aquarium filled with 5 L of fresh ASW (50 × 20 × 20 cm), covered with a dark curtain, and acclimated to the test environment (>1 h). Then, 5 mL of vehicle or 5,6,11-trideoxyTTX (10^−5^ M) was administered to one side of the aquarium (the final estimated concentration of 5,6,11-trideoxy TTX was 10^−8^ M). Before administration, the fish intermittently swam left and right and occasionally stopped in one place in the aquarium. Administration of 5,6,11-trideoxyTTX caused the fish to approach and frequently stop on the administered side (**Movie S1**). Such change in the swimming pattern was reflected in the time vs. x-position plot of the fish (**Fig. 4a**). Similar changes in swimming patterns were observed with food extract administration (Supplementary **Fig. S1**), but not with vehicle administration (**Fig. 4d**).

**Figure 4:**
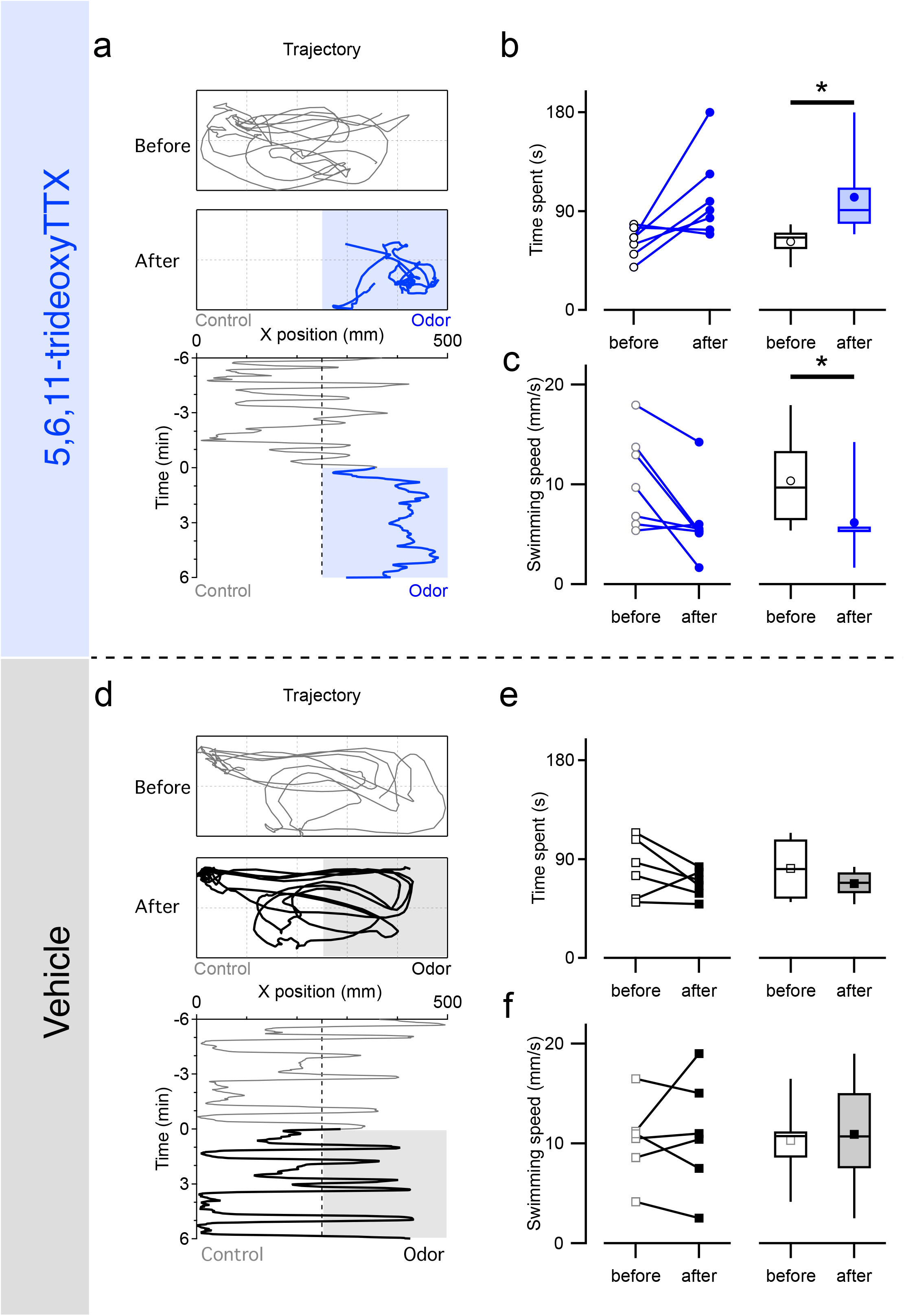
5,6,11-trideoxyTTX attracts grass puffers. Tracked location of a grass puffer before and after 5,6,11-trideoxyTTX administration (5 mL of 10^−5^ M dissolved in 2.5 × 10^−4^ N acetic acid solution, Glid: 100 mm) and positions of the fish are plotted along the horizontal axis of the aquarium every 1 s. 5,6,11-trideoxyTTX was applied at time 0 and remained in the aquarium thereafter (**a**). Time spent on the administered side (**b**) and mean swimming velocity (**c**) during the 3-minute periods before and after the 5,6,11-trideoxyTTX administration (n=7). Paired *t*-test compared before and after the administrations. (Time spent, P = 0.037). *p < 0.05. Tracked location of a grass puffer before and after vehicle administration (5 mL of 2.5 × 10^−4^ N acetic acid solution, Glid: 100 mm) and positions of the fish are plotted along the horizontal axis of the aquarium every 1 s. Vehicle was applied at time 0 and remained in the aquarium thereafter (**d**). No significant changes in the time spent (3-min period) (**e**) or swimming velocity (mm/s; **f**) in response to the vehicle administered on one side of the experimental aquarium (n=6). Paired *t*-test compared before and after the administrations. (Time spent, P = 0.202; Swimming velocity, P=0.728). Open or colored dots and boxes in the plot represent the data before and after odorant administrations. The line and scatter plots show the value from individual fish, and box plots show the median, quartiles (boxes), and 10%–90% range (whiskers).

To analyze odorant preference, we calculated the time spent on the administered side and the mean swimming velocity of the individual grass puffer during the three-minute periods from the second to the fifth minute before and after the administration, when the odorant were considered to have sufficiently diffused to the administered side in the aquarium. This period was determined based on the preliminary dye diffusion test (see Materials and Methods; excluded the data within 2 minutes before and after the start of odorant administration from the analysis to exclude the effects associated with the administration procedure). The time spent by the grass puffers on the administered side increased (62.1 ± 5.1 to 102.9 ± 14.6 s, *p*<0.05; **Fig. 4b**), and the swimming velocity of grass puffers decreased after the 5,6,11-trideoxyTTX administration (10.4 ± 1.8 to 6.2 ± 1.5 mm/s, n=7, *p*<0.05; **Fig. 4c**). In contrast, there are no significant changes in time spent (81.5 ± 10.8 to 67.7 ± 5.1 s; **Fig. 4d**) and swimming velocity (10.3 ± 1.6 to 10.9 ± 2.3 mm/s; **Fig. 4e**) during the three-minute periods before and after the vehicle administration (n= 6 fish). Furthermore, we examined the behavioral response of non-TTX bearing fish to 5,6,11-trideoxyTTX using medaka (*Oryzias latipes*; inhabiting freshwater without any TTX-bearing prey organisms; Supplementary **Fig. S2**). Medaka did not prefer to stay on the administered side of 5,6,11-trideoxyTTX. These results suggest that grass puffers prefer 5,6,11-trideoxyTTX.

### Activity-dependent labeling of 5,6,11-trideoxyTTX sensitive OSNs

To identify OSN that responded to 5,6,11-trideoxyTTX, we first observed the gross morphology of the OE from the grass puffer. Differential interference microscopy images of live olfactory lamellae showed that the surface of olfactory lamella had repeated structures in which the elliptical “islets” of the sensory epithelium (**se** in **Fig. 5a)** were bordered by oval cells and exposed on the surface (arrowheads in **Fig. 5a**). The cross-sectional structure of the olfactory lamellae showed that oval cells (arrowheads in **Fig. 5b**) surrounded small islets where there were some cilia on the surface (arrow in **Fig. 5b**).

**Figure 5:**
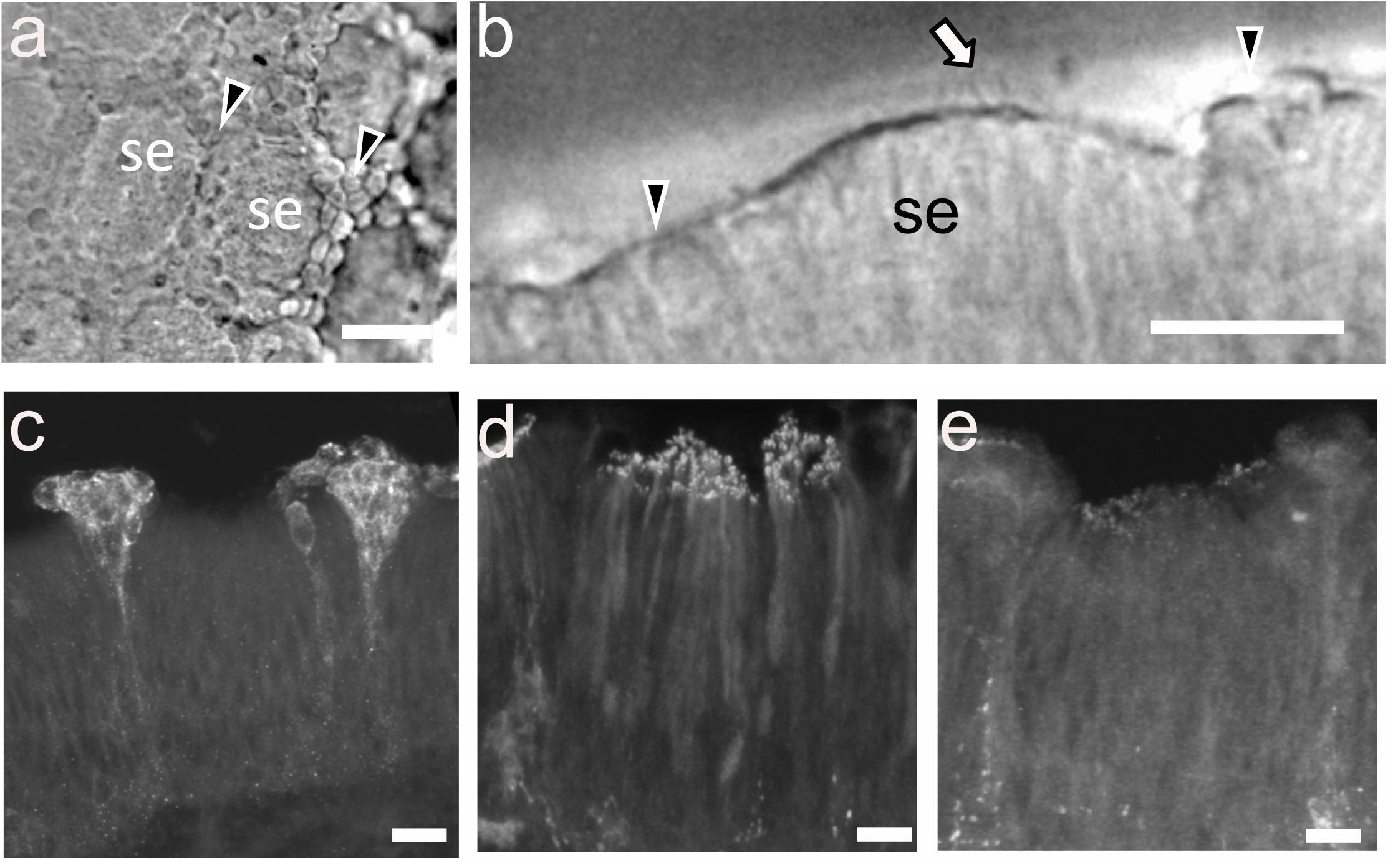
Activity-dependent labeling by pERK antibody labeled oval-shaped cells on the surface of the olfactory epithelium. Differential interference contrast photomicrographs of the surface (**a**) and the edge (**b**) of the olfactory lamella of grass puffer. Arrowheads represent the non-ciliated epithelium surrounding the sensory epithelium “islets” (SE). Arrow represents sensory cilia on the surface of the sensory epithelium. Scale bar = 20 μm. pERK-immunopositive OSNs after exposure to 5,6,11-trideoxyTTX (**c**) or L-Arg (**d**). No labeled cells were observed with the Vehicle control (**e**). Scale bar: 5 μm. Six fish examined for each stimulus showed similar results.

We performed immunohistochemistry of excised OE from grass puffers with the neural activity marker (phosphorylated extracellular signal regulated kinase; pERK) antibody as an activity indicator. Administration of 5,6,11-trideoxyTTX (10^−6^ M for 1min) labeled oval cells clustering on the surface of the OE (**Fig. 5c**), whereas L-Arg administration (10^−6^ M for 1min) labeled elongated cells with strongly labeled cilia whose somata were in the middle layer of the OEs (close to the basal membrane; **Fig. 5d**). In addition, administration of 5,6,11-trideoxyTTX also induced an activity-dependent uptake of Alexa Fluor 555-conjugated dextran (10 kDa) to oval cells in the surrounding region of the islets (Supplementary **Fig. S3**). However, vehicle administration did not label any OSNs (**Fig. 5e**).

## Discussions

In this paper, we electrophysiologically, morphologically, and behaviorally demonstrated that one of the TTX analogs, 5,6,11-trideoxyTTX, is an odorant for grass puffers, *T. alboplumbeus*. A previous behavioral study reported that TTX attracts male grass puffers during the spawning period ^10^. TTX also attracted juvenile tiger puffers regardless of sex ^11–13^, and this attraction effect was blocked by nose ablation ^11^. However, TTX did not induce any EOG responses in our study. Previous studies used crude TTX extracted from the wild-caught tiger puffer ovary or liver ^11^. The original TTX extraction method used in previous studies reported that their crude TTX from pear puffer (*T. vermicularis*) ovaries contained both TTX and TTX analogs, including 5,6,11-trideoxyTTX, 4-*epi* TTX, and 4,9-anhydroTTX ^14^. However, our electrospray ionization mass spectrometry analysis indicated that the commercially available TTX that we used in our study did not contain detectable levels of these TTX analogs (data not shown; except 4,9-anhydroTTX because both 4,9-anhydroTTX and TTX are epimerized each other in aqueous solution and are in chemical equilibrium.). Therefore, since the crude TTX extracts obtained in previous studies contain TTX analogs, we hypothesized that some of these TTX analogs might act as odorants for grass puffers and examined the effects of two types of synthesized TTX analogs (4,9-anhydroTTX and 5,6,11-trideoxyTTX). In our recordings, 5,6,11-trideoxyTTX induced an EOG response in grass puffers, but 4,9-anhydroTTX and TTX did not. In addition, the EOG amplitude of 10^−5^ M 5,6,11-trideoxyTTX was significantly greater than that at the same concentration of L-Arg, which is a well-known odorant for many teleosts (Figure 3a). Furthermore, 5,6,11-trideoxyTTX induced EOG responses even at 10^−8^ M (Figure 3 c and d). Because the stimulation threshold of amino acids on the fish olfactory system is 10^−9^ M to 10^−7^ M ^15^, 5,6,11-trideoxyTTX is considered to be fully capable of being an odorant for grass puffers.

5,6,11-trideoxyTTX elicits chemotaxis of grass puffers. In response to 5,6,11-trideoxyTTX administration to one side of the aquarium, grass puffers were attracted to the administered side and showed the behavior of slowing down and swimming in that area. Our preliminary dye diffusion test estimated that the grass puffers could recognize 5,6,11-trideoxyTTX ranging 2 × 10^−8^ to 1 × 10^−5^ M within 2 minutes after administration (see Materials and Methods). These estimated 5,6,11-trideoxyTTX concentrations correspond to the threshold concentration of the EOG response, although the concentration is much higher than those of the previous behavioral studies. Mature male grass puffers were attracted to 1.5 × 10^−11^ M crude TTX extract ^10^. Male and female juvenile tiger puffers could also recognize 3.7 × 10^−9^ ^13^ and 2.6 × 10^−8^ M ^11^ of crude TTX extract, respectively. In these studies, they started their “choice” experiments after diffusion of TTX from agar or gelatin on one side of the Y-maze aquarium. They then examined which side of the Y-maze aquarium the pufferfish swam in. Alternatively, TTX analogs other than 5,6,11-trideoxyTTX may also be involved in chemoreception synergistically. For example, female sex pheromones in moths usually consist of multicomponent blends of five to six hydrocarbons, including unbranched fatty acids, alcohols, acetates, or aldehydes in particular combinations and ratios ^16^. These differences seem to contribute to effective TTX concentrations in previous studies and that of 5,6,11-trideoxyTTX in our study.

Immunohistochemistry with pERK antibody suggested that 5,6,11-trideoxyTTX activates oval cells distributed in the border area around the ciliary “islets” on the olfactory lamella, which were classified as undifferentiated epithelium in previous study ^17^. These oval cells were also labeled with simultaneous administration of fluorescent dextran with 5,6,11-trideoxyTTX. It is widely known that teleost fish are equipped with three types (ciliated, microvillous, and crypt) of OSNs intermingled in the OE but show distinct morphology ^18^. Among them, crypt OSNs have an oval shape, are located close to the epithelial surface, and have few cilia and microvilli ^19^. Cells labeled by 5,6,11-trideoxyTTX co-administration had no cilia or microvilli and were located on the epithelial surface around the islets of ciliated epithelium. Based on these morphological characteristics, the 5,6,11-trideoxyTTX-sensitive OSNs are presumed to be crypt or another unknown type of OSN. Our recent study using the green spotted puffers suggested that cells immunopositive for S-100, a marker protein of crypt-type OSNs, were also labeled by another neuronal activity marker, phosphorylated ribosomal protein S6, antibody that is formed after 5,6,11-trideoxyTTX administration ^20^. This result further supports that the 5,6,11-trideoxyTTX sensitive OSNs are crypt-type OSNs.

Here, let us consider the physiological significance of 5,6,11-trideoxyTTX as an odorant for the grass puffers. 5,6,11-trideoxyTTX has been reported to be present in the body of toxic pufferfish. The distribution of TTX and its analogs throughout the body of grass puffers was reported, and 5,6,11-trideoxyTTX accumulated mainly in the ovary ^21^. A receptor binding assay study of voltage-gated sodium channels in rat brain suspensions reported that the dissociation equilibrium constant of 5,6,11-trideoxyTTX (> 5000 nM) was extremely high compared with that of TTX (1.8 ± 0.1 nM) ^22^. With such a low affinity for voltage-gated sodium channels, 5,6,11-trideoxyTTX can hardly act as a toxin, suggesting that it cannot function as a defense substance against predators. Therefore, the biological significance of why toxic pufferfish accumulate such almost nontoxic TTX analog remains unclear.

One possibility is that 5,6,11-trideoxyTTX functions as an attractive sex pheromone for toxic pufferfish. It has been reported that TTX is secreted from the secretory glands on the skin of *T. pardalis, T. flavipterus, T. vermicularis*, and grass puffers ^6^ and is released from ovulated oocytes in the grass puffers ^10^. Interestingly, grass puffer’s whole body TTX content was higher during the spawning period ^23^. Because 5,6,11-trideoxyTTX is particularly abundant in pufferfish ovaries ^24^, 5,6,11-trideoxyTTX may also be leaked from the ovarian luminal fluid with TTX during spawning periods, which attracts the males.

Another possibility is that grass puffers use 5,6,11-trideoxyTTX as an olfactory cue to locate and feed on TTX-bearing organisms. Wild toxic puffers tend to feed on benthic organisms, such as flatworms, starfish, small gastropods, and skeleton shrimp that are known to possess TTX ^2^. For example, a recent study reported that a TTX-bearing planocerid flatworm was found in the gut content of grass puffers ^25^ and *Chelondon patoca* ^26^. Furthermore, some carnivorous gastropods have 5,6,11-trideoxyTTX in addition to TTX (*Charonia lampas lampas*^27^; *Nassarius spp.* ^28^), and toxic snails have been reported to be attracted to crude TTX extracts ^29^. Additionally, grass puffers ingested toxic eggs of another pufferfish, *T. pardalis,* which contained 5,6,11-trideoxyTTX ^30^. All these studies support the idea that the pufferfish seems to be attracted by 5,6,11-trideoxyTTX and feed on TTX-bearing organisms, resulting in TTX accumulation.

In conclusion, the grass puffers detect 5,6,11-trideoxyTTX as an odorant. Our findings suggest that pufferfish sense a TTX analog, which is present in their body with TTX but whose function was unknown, as an odorant. Further studies are needed to determine whether other TTX analogs can also be detected by olfaction and whether multiple TTX analogs synergistically induce such effects. However, we believe that our results provide a new perspective on the use of TTX and its analogs by toxic puffers.

## Materials and Methods

### Animals

One hundred seventy-four grass puffers (body length, 5–12 cm; bodyweight, 6–32 g) were collected from the coastal waters of Ise-bay in Minamichita-Cho on the Pacific coast of central Japan by line-fishing. Grass puffers were fed commercial diets (Tetra Krill-E; Spectrum Brands, Madison, WI, USA) in aerated 300 L breeding tanks (10–25°C, approximately 50 fish per tank) filled with ASW (Instant Oceans Sea Salt; Instant Ocean, Blacksburg, VA, USA) under a 12-h light/12-h dark photoperiod. In this study, we used randomly selected fish from the breeding tank in the experiment. Thus, both males and females were used without distinction. Animal Use and Care Protocols were approved by the Center for Animal Research and Education of the Nagoya University (Approved number: A210871-004). All animals were maintained and used in accordance with ARRIVE guidelines and the Tokai National Higher Education and Research System Regulations on Animal Care and Use in Research (Regulation No.74). No statistical methods were used to predetermine the sample size because many of the outcomes were unknown.

### Odorant chemicals

TTX (for biochemistry, citrate buffered, FUJIFILM Wako Pure Chemical Corporation, Osaka, Japan) was dissolved in ultra-pure water at 1.5 × 10^−3^ M. Two types of TTX analogs, 4,9-anhydroTTX and 5,6,11-trideoxyTTX, were chemically synthesized and purified by HPLC on ion-exchange column (Hitachi gel 3013C). These synthesized analogs do not contain other derivatives and reagents before use, judging by their ^1^H and ^13^C-NMR spectra. These purification procedures and the spectra were described in the papers and the supporting information cited in references ^31^ and ^32^, respectively. Synthesized TTX analogs were dissolved in 2.5 × 10^−2^ N acetic acid in artificial cerebrospinal fluid (ACSF; NaCl 231 mM, KCl 8.1 mM, MgCl_2_ 1.3 mM, CaCl_2_ 2.4 mM, D-glucose 10 mM, 4-(2-hydroxyethyl)-1-piperazineethanesulfonic acid (HEPES) 10 mM; pH adjusted 7.4 with NaOH), that was modified from the Ringer’s solution for marine fish^33^, at 10^−3^ M and stored at −30°C until use. L-Arg (FUJIFILM Wako) was prepared as a 10^−2^ M stock in ACSF. For testing, test odorant solutions were prepared fresh as required by diluting aliquots of concentrated stock in ASW. L-Arg was always tested at 10^−5^ M. When testing a series of solutions with different concentrations, the lowest dose was tested first, followed by solutions with concentrations that increased by a factor of 10 to 10^−4^ M. A vehicle solution was prepared in an identical manner by diluting 2.5 × 10^−2^ N acetic acid in ASW (final concentration, 2.5 × 10^−4^ N). The food extract solution was prepared by obtaining the supernatant of ground commercial diet (0.5 g) suspended in ASW (50 mL).

### Electroolfactogram recordings

EOG recordings were performed on secured grass puffers on a holding apparatus that was placed into a plastic experimental trough. The tricaine methanesulfonate (MS–222; 0.02%) anesthetized grass puffer was mounted on the center groove of a hand-made plastic holder by wrapping it in a paper towel and aluminum foil after attaching an Ag–AgCl plate (1 × 1 cm) with the body wall as a reference electrode. The gills were perfused with ASW containing 0.01% MS–222 through a tube inserted into the mouth throughout the experiments. The skin and cartilage covering the right olfactory cavity were opened, and the exposed olfactory cavity was continuously perfused with ASW using an 18 G stainless needle (NN–1838S; Terumo, Tokyo, Japan). The perfusion rate of ASW was 1.0 mL/min. A computer-controlled solenoid valve (UMG1–T1; CKD, Komaki, Japan) delivered 10-s pulses of test solutions by switching between the ASW and test solutions (diluted appropriately in ASW) at 3-min intervals.

EOG responses were recorded using a saline–agar (0.6%)-filled glass capillary (tip diameter, 100 μm) that was bridged with an Ag–AgCl wire filled with ASW. The indifference electrode was placed lightly on the head skin surface near the perfused olfactory cavity, and the recording electrode was positioned over the olfactory rosette using micromanipulators. The electrodes were insulated with paraffin–Vaseline paste that was placed on the surface of the fish’s head. Electrical signals were differentially amplified (gain 1,000×; Band-pass 0.08–100 Hz) using a DC preamplifier (AVH-11 in VC-11 digital oscilloscope; Nihon-Kohden, Tokyo, Japan). In this study, recordings were not made in full DC mode to minimize the slow drift of the baseline caused by the changing contact area between the odorant solutions constantly dripped into the nose and the artificial seawater being perfused on the gill for respiration. The amplified EOG signal was digitized (10 kHz) and stored on a laptop computer using a USB-6251 data acquisition device (National Instruments, Austin, TX, USA) and WinEDR software (ver. 3.2.7; http://spider.science.strath.ac.uk/sipbs/software_ses.htm; Strathclyde University, Glasgow, UK).

### Behavioral experiments

Behavioral experiments were performed using adult male and female grass puffers. For the behavioral assay, a grass puffer was transferred to a test aquarium (50 × 20 × 20 cm) filled with 5 L of fresh ASW. The test aquarium was covered with a black curtain and illuminated by an infrared LED-array illuminator (940 nm, ca. 2 W; AE-LED56V2, Akizuki Denshi Tsusho Co., Tokyo, Japan) so that the experimenter was invisible to the fish. A video camera (XC-ST50, Sony, Tokyo, Japan) equipped with a low distortion C-mount lens (f = 4.4 mm; LM4NC1M, Kowa, Nagoya, Japan) was mounted vertically above the aquarium, and the camera system was adjusted so that the aquarium filled the camera frame. The fish was allowed to acclimate to a dark environment for 90–180 min.

Time-lapse image acquisitions from the top of the test aquarium were started (1 s intervals). The time-lapse images were acquired on a laptop computer using image acquisition software (Micro-Manager ver. 1.4.22; https://micro-manager.org)^34^ and a USB-video capture unit (PCA-DAV, Princeton, Tokyo, Japan) at a resolution of 640 × 480 pixels. After 6 min of image acquisition, 5 mL of 10^−5^ M 5,6,11-trideoxyTTX or vehicle (5 mL of 2.5 × 10^−4^ N acetic acid) was gently applied to one side of the test aquarium through a silicone tube attached to a 10 mL syringe. We randomly altered the administered side of odor solutions. Then, image acquisitions were continued for up to 6 min. We excluded the data that fish stopped swimming longer than 5 seconds before the odorant administration (20% of the experiments). A preliminary test indicated that diffusion indicator dye (1% methylene blue in ASW) reached half of the experimental aquarium where a grass puffer was swimming within 2 min after administration and had spread throughout the aquarium at the end of the observation period. The numeric x–y coordinates representing the movement of the fish over time (one coordinate in a frame) were extracted from each video using the UMA tracker software (Release-15; http://ymnk13.github.io/UMATracker/)^35^.

### Activity-dependent labeling of OSNs

To study the ligand specificity of OSNs, we excised the olfactory lamellae from anesthetized fish. The excised olfactory lamella was washed three times for 5 min in ASW and stimulated with odorant solutions (vehicle, L-Arg, TTX, or 5,6,11-trideoxyTTX; 10^−6^ M, 1 mL, diluted with ASW) for 5 min in a 1.5 mL tube. After odor stimulation, OE was fixed in 4% paraformaldehyde and 15% picric acid in 0.1 M PBS overnight at 4°C, rinsed in PBS, and cryoprotected in 30% sucrose in PBS overnight at 4ºC. Cross cryosections (14–20 μm) of OE mounted onto gelatin-coated slides and dried. After rehydration in PBS, the sections were blocked in 2% normal goat serum (NGS) in PBS with 0.05% Triton-X100 (PBST) for 3 h at 28°C. The sections were then incubated with anti-phosphorylated-p44/42 mitogen-activated protein kinase (MAPK; extracellular regulated kinase, ERK1/2) rabbit monoclonal antibody (#4370, Cell Signaling Technology, Danvers, MA, USA; 1:500), which was used in zebrafish OE^36^, as a neural activity marker that was diluted with 0.2% NGS in PBST for 2 days at 4°C. The sections were washed three times for 5 min with 0.2% NGS in PBST. Sections were then incubated with Alexa Fluor 488-conjugated secondary antibody (goat anti-rabbit IgG (H+L); A-11008, Thermo Fisher Scientific, 1:500) diluted with 0.2% NGS in PBST for 1 day at 4°C.

We also used odor-induced uptake of fluorescent dye-conjugated dextran ^37^. The fish was anesthetized using 0.02% MS-222 and secured on the holding apparatus used in EOG experiments. The gills were continuously perfused with ASW containing 0.01% MS-222 through a tube inserted into the mouth throughout the procedure. The skin and cartilage covering the olfactory cavity were opened, and the exposed olfactory cavity was continuously perfused with ASW using an 18G stainless needle. The olfactory cavity of anesthetized fish was perfused with L-Arg (10^−5^ M) or 5,6,11-trideoxyTTX (10^−5^ M) together with a dextran conjugated with Alexa Fluor 555 (M.W. 10,000, 10^−6^ M; D34679, Thermo Fisher Scientific, MA, USA) for 5 min. In the control experiments, the olfactory cavity of anesthetized fish was exposed to only Alexa Fluor 555-dextran (10^−6^ M) for 5 min. The flow rate was approximately 1.0 mL/min. After the odorant application, the olfactory cavity was rinsed with a continuous flow of ASW for at least 5 min.

Fish were kept alive for 3 to 4 h in aerated ASW at room temperature. Then, the fish was reanesthetized using 0.1% MS-222 and decapitated. The decapitated head was immersed in 4% paraformaldehyde in 0.1 M PBS (pH 7.2, 4°C, 1 h). After fixation, the olfactory lamellae were dissected and rinsed with PBS. The olfactory lamellae were then counterstained with 4ʹ,6-diamidino-2-phenylindole (DAPI; 1 ng/mL, 5 min) and mounted on a hole glass slide with mounting medium (10% Mowiol 4-88, 25% glycerol, 0.2 M Tris-HCl (pH 9.0), and 2.5%-3.0% 1,4-diazabicyclo [2,2,2] octane; diluted with ultrapure water).

A confocal microscope (FluoVIEW 1000D, IX81, Olympus, Tokyo, Japan) was used to observe the details of olfactory lamellae. Pictures were taken in planes and were separated by 0.45 μm z-axis steps. Confocal microscopy images were analyzed using FluoRender (ver. 2.26.3; http://www.sci.utah.edu/software/fluorender.html) for three-dimensional (3D) reconstruction and visualization of dextran-labeled cells^38^. Pre-processing of images before 3D reconstruction (median filtering [radius: 1 pixel] and trimming) was performed by Fiji (https://fiji.sc/)^39^.

### Data analysis

All animal experiments were performed on fish of both sexes. Statistical analyses and graph preparations in this study were performed using R (ver. 4.0.4; https://www.r-project.org) and Igor Pro (ver. 9.00; WaveMetrics Inc., Lake Oswego, OR, USA). Data are presented as a dot plot or a box–whisker plot (box: median and quartiles; whiskers: 10%–90% range) or mean ± standard error of the mean (SEM).

## Acknowledgment

We would like to express our gratitude to Prof. Naoyuki Yamamoto for his advice during the experimentation.

## Author contributions

Y.N. and H.A. conceived the study, designed experiments, and wrote the manuscript. R.S., R.N., M.A., and T.N. synthesized and purified TTX analogs. Y.N. performed electrophysiology, behavioral experiments, and activity labeling of olfactory sensory neurons by fluorescent dextran. T.S. and K.M. performed immunohistochemistry of olfactory sensory neurons.

## Data availability

The datasets generated during the current study are available from the corresponding author on reasonable request.

