## Supplemental Data 1 for "An almost nontoxic tetrodotoxin analog, 5,6,11-trideoxytetrodotoxin, as an odorant for the grass puffer"

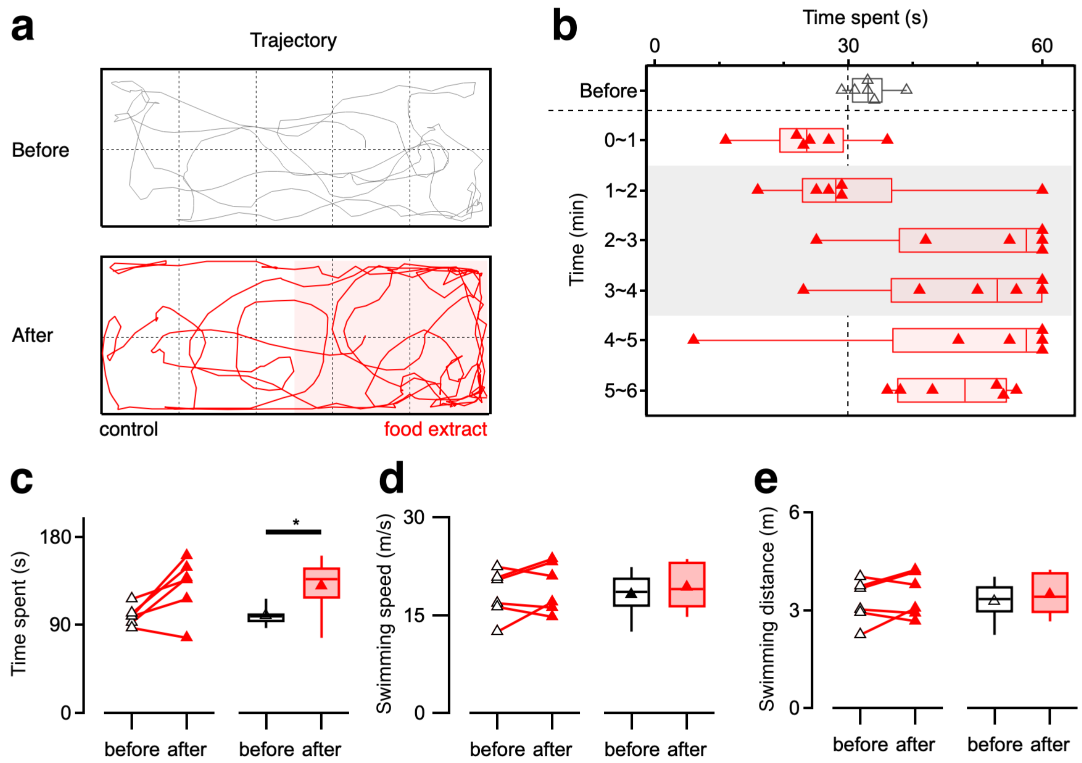


**Figure S1: Food extracts attracts grass puffers.**

Tracked location of a grass puffer before and after food extract administration (5 mL of supernatant of 0.5 g ground commercial diets suspension in 50 mL artificial sea water, Glid: 100 mm; **a**). Changes in the time (1-min period) spent by grass puffers in response to food extracts administered on one side of the experimental aquarium(n=7; **b**). The vertical dotted line represents the chance level (= No odorant preference). Open or colored triangles and boxes in the plot represent the data before and after the food extracts administrations. Dot plots show the value from individual fish, and box plots show the mean (markers), median, quartiles (boxes), and 10%–90% range (whiskers). Time spent on the administered side (**c**), swimming speed (**d**), and swimming distance (**e**) during the 3-minute periods before and after the food extract administration. The line and scatter plot in **c** ~ **e** show the changes of individual puffers before and after the administration, and the box plots show the mean (markers), median, quartiles (boxes), and 10% - 90% range (whiskers). Paired *t*-test compared before and after the administrations. (Time spent, P = 0.037). $^*p < 0.05$.

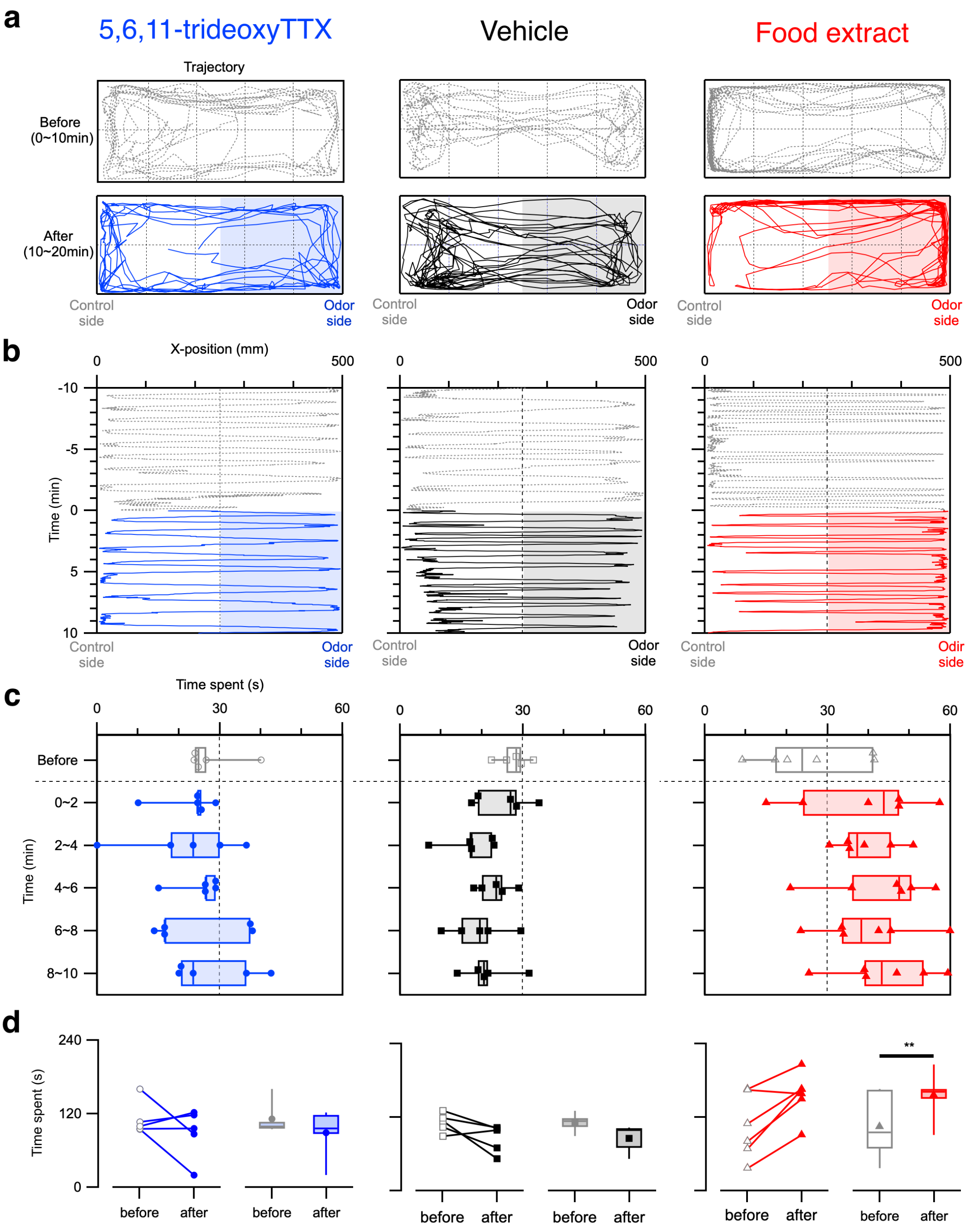


**Figure S2: 5,6,11-trideoxy TTX did not attracts nonTTX-bearing fish.**

(**a**) Tracked location of a medaka, a nonTTX-bearing fish, before and after 5,6,11-trideoxyTTX (left), vehicle (middle), or food extracts administration (right). Glid: 100 mm. (**b**) Positions of the fish are plotted along the horizontal axis of the aquarium every 1 s. Test solutions were applied at time 0 and remained in the aquarium thereafter. (**c**) Changes in the time spent (2-min period) by medaka in response to test solutions administered on one side of the experimental aquarium (n = 5 for 5,6,11-trideoxyTTX and vehicle; n = 6 for food extracts). The vertical dotted line represents the chance level (= No odorant preference). Open or colored dots and boxes in the plot represent the data before and after the test solutions administrations. Dot plots show the value from individual fish, and box plots show the median, quartiles (boxes), and 10%–90% range (whiskers). (**d**) Time spent on the administered side during the 4-minute periods before and after the test solutions administrations. The line and scatter plot show the changes of individual puffers before and after the administration, and the box plots show the mean (markers), median, quartiles (boxes), and 10% - 90% range (whiskers). Paired *t*-test compared before and after the administrations (food extract, P=0.014). ** $p < 0.01$.

dr-R strain of medaka (*Oryzias latipes*) were used in this experiment. Eight to Sixteen fish of both female and male were maintained in a fish tank with water circulation under 14-h light/10-h dark photoperiod (light on at 09:00 and off at 23:00) condition at a water temperature of 27 ± 2 °C. These were fed three or four times per day with live brine shrimp and/or commercial diets (Tetra Medaka-bijin; Spectrum Brands, Yokohama, Japan). The experimental procedure was the same as for the grass puffers except for the lighting condition, the use of deionized freshwater (3 L) instead of artificial seawater (5 L), and the image acquisition durations of post odorant administrations.

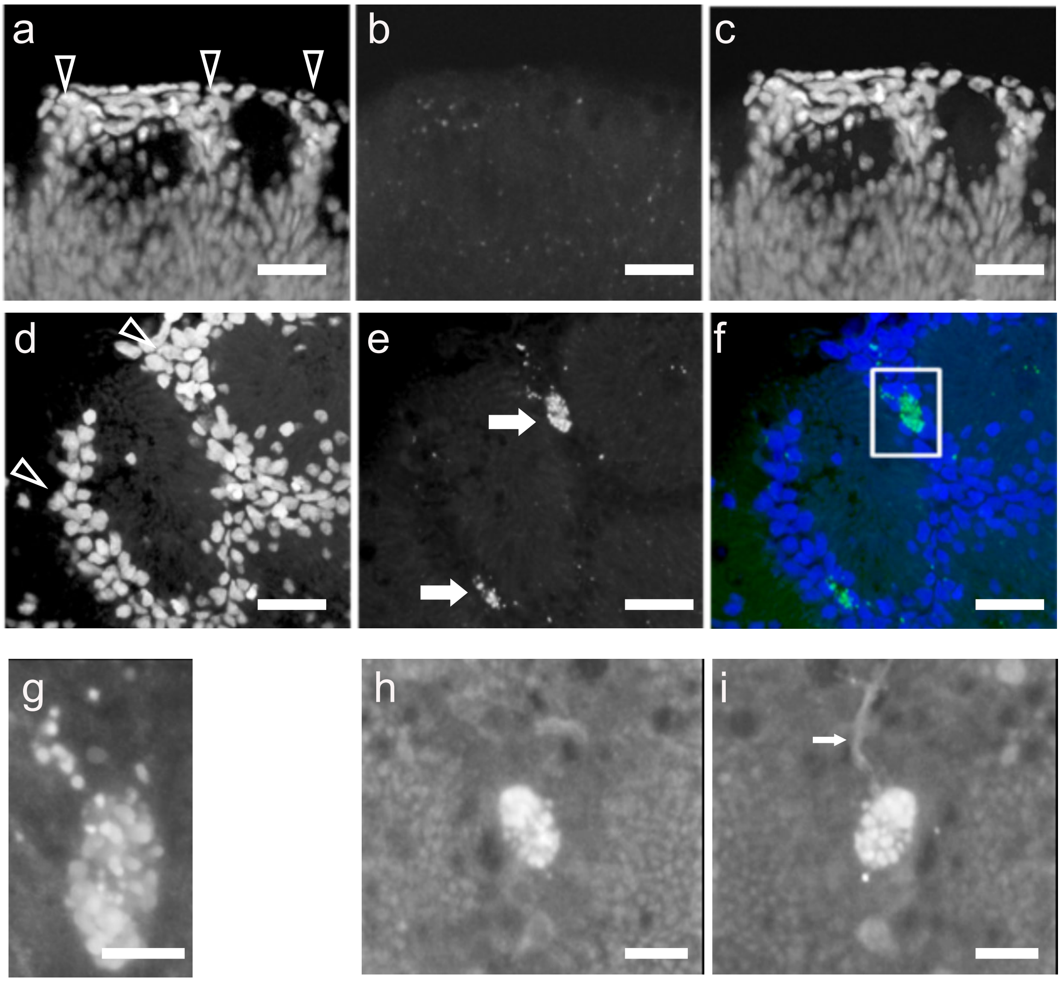


**Figure S3: Activity-dependent fluorescent dextran uptake labeled 5,6,11-trideoxyTTX responding olfactory sensory neurons on the olfactory epithelium of the grass puffer.**

Confocal microscopy images of the olfactory lamella after vehicle administration (**a–c**). Nuclear staining using DAPI (**a**) visualized the cells surrounding the islets whose nuclei were located on the surface of the olfactory lamella. Co-administration of Alexa Fluor555-dextran and vehicle did not label any cell-like structures (**b**). Superimposed image of **a** and **b** (**c**). The image is a z-projection of 41 pictures (z-step, 0.45 μm).

Confocal microscopy image of the olfactory lamella after 5,6,11-trideoxyTTX administration (**d–f**). Nuclear staining using DAPI (**d**), Alexa Fluor555-dextran (**e**), and their composite image (**f**). Co-administration of Alexa Fluor555-dextran labeled the large oval cells that were in the surrounding region of the islets (arrows in **e**). Scale bar = 20 μm. The image is a z-projection of 28 pictures (z-step, 0.45 μm).

Enlarged image of the rectangular region in **f** (**g**). Alexa Fluor 555 labeling was frequently seen as bright granular aggregations within the cell. Scale bar = 5 μm.

Another z-projection image (17 planes with a z-step of 0.48 μm) of Alexa Fluor555-dextran labeled cells by 5,6,11-trideoxyTTX administration (surface side view, **h**; and basal membrane side view, **i**). The arrow indicates an axon-like structure that extends from the Alexa Fluor555-dextran-labeled cell. Scale bar = 5 μm. Five fish examined for each stimulus showed similar results.

**Movie S1: Attractive swimming behavior of grass puffers toward 5,6,11-trideoxyTTX observed under infrared illumination.**

Vehicle or 5,6,11-trideoxyTTX solutions were added from the right side of the test aquarium 6 minutes after the start of image acquisitions.
